# Novel synergistic combinations of last-line antibiotics and FDA-approved drugs against Klebsiella pneumoniae revealed by in vitro synergy screenings

**DOI:** 10.1101/2022.05.16.491802

**Authors:** Marta Gómara-Lomero, José Antonio Aínsa, Santiago Ramón-García

## Abstract

Treatment of infections caused by multi-drug resistant (MDR) enterobacteria remains challenging due to the limited therapeutic options. Drug repurposing could accelerate the development of urgently needed successful interventions. This work aimed to identify and characterize novel drug combinations against *Klebsiella pneumoniae* based on the concepts of synergy and drug repurposing. We performed a semi-qualitative high-throughput synergy screening (sHTSS) with tigecycline, colistin and fosfomycin (last-line antibiotics against MDR Enterobacteriaceae) combined with an FDA-library containing 1,430 clinically approved drugs. Selected hits were further validated by secondary checkerboard (CBA) and time-kill (TKA) assays. Our sHTSS results yielded 37, 31 and 41 hits showing synergy with tigecycline, colistin and fosfomycin, respectively. Most hits (75%) were known antibiotics. Non-antibiotic compounds included other anti-infective agents (7%), antineoplastics (7%) or antipsychotics (3%). Overall, 15.09% and 65.85% of hits were further confirmed by CBA and TKA, respectively, indicating that TKA is more useful than CBA for the validation of synergistic combinations. Accordingly, TKA were used for synergy classification based on determination of the bactericidal activities at 8, 24 and 48 hours. Twenty-seven combinations were validated with effective synergistic activity against *K. pneumoniae* by TKA, six of them novel non-antibiotic combinations. Based on our observations we conclude that repurposing approaches allowed to enhance the activity of last-line antibiotics in the treatment of MDR *K. pneumoniae*. sHTSS paired to TKA was a powerful tool for the identification of novel synergistic drug combinations against *K. pneumoniae*. Further pre-clinical studies might support the translational potential of these novel combinations.

## INTRODUCTION

Antimicrobial resistance (AMR) is a major global public health problem. There were an estimated 4,95 million deaths associated with bacterial AMR in 2019, including 1,27 million deaths directly attributable to bacterial AMR (1). In Europe, AMR is responsible each year for about 33,000 deaths and 1,1 billion Euros of costs to EU/EEA health care systems (2). Carbapenemase-producing *Klebsiella pneumoniae* causes severe infections associated with 23% to 75% mortality rates (3); it is considered among the most concerning pathogens with prevalence of carbapenem resistance worldwide increasing more than seven-fold in the EU since 2006. In addition, resistance to two or more antimicrobial groups is growing concern (4,5).

The acquisition of resistance to most available antibiotics severely limits therapeutic options of infections caused by MDR Enterobacteriaceae. Clinicians are thus forced to use last-line antibiotics in combination therapy with mixed success rates (6,7). In 2017, WHO highlighted drug-resistant enterobacteria as priority pathogens for which new antibiotics were urgently required (8); however, the traditional process of drug discovery and development is long and costly (9,10). Although new clinical options have been recently approved, scarce evidence and lack of randomized clinical trials prevent from establishing optimal clinical guidelines. New strategies changing the current paradigm are thus urgently needed (11).

Drug repurposing has been largely regarded as a promising alternative to the traditional drug discovery and development process (12,13), since safety profiles for a new indication could be implemented faster as long as they are used following originally approved recommendations. For example, the antifungal amphotericin B was repositioned for the treatment of visceral leishmaniasis, a parasitic infection (14). In addition, the search of synergistic partners might increase, on the one hand, the therapeutic range of drugs whose potential use could be limited due to toxicity issues and, on the other hand, rescue antimicrobials not reaching efficacy breakpoints (15,16).

In this study, we aimed to identify partners that could potentiate the activity of three last-line antibiotics used against MDR *K. pneumoniae*. For this, we performed a semi high-throughput synergy screening (sHTSS) of an FDA-approved drug library in combination with tigecycline, colistin and fosfomycin (17). Lead combinations were validated by checkerboard (CBA) and time-kill (TKA) assays. Synergistic and killing effects were evaluated at different time points to obtain a priority list of combinations with potential for clinical translation.

## MATERIALS AND METHODS

### Bacterial strains, media and chemicals

*Klebsiella pneumoniae* ATCC 13883 was used for all experiments. Bacterial stocks (15% glycerol) were preserved at −20°C in LB broth. A new stock was thawed for every experiment to ensure assay robustness, and sub-cultured on Mueller Hinton broth (MHB) for 24 hours before each MIC, CBA and TKA assay were performed in CAMHB. Drug susceptibility testing in solid media and sHTSS were performed in Mueller-Hinton agar (MHA). All cultures were incubated at 36-37°C.

The FDA-approved drug library was purchased from Selleckchem (catalogue #L1300). Drugs were purchased from the European Pharmacopoeia (Strasbourg, France), except fosfomycin. sHTSS FDA-hits were obtained from Sigma–Aldrich (Darmstadt, Germany). Drugs were freshly reconstituted in DMSO or water.

### Drug susceptibility testing

MIC determinations were performed by broth microdilution according to CLSI guidelines (18) and the MTT [3-(4,5-dimethylthiazol-2-yl)-2,5-diphenyl tetrazolium bromide] assay (17,19). Briefly, two-fold serial dilutions were inoculated with 5×10^5^ CFU/mL (OD_550_=0.2 corresponded to 1.5×10^8^ cell/mL) in 96-well plates (V_F_= 150 μL/well) and incubated for 18-20 hours. After incubation, 30 μL/well of 5 mg/mL MTT plus 20% Tween 80 were added and incubated for 3 hours. MIC values were defined as the lowest concentration of drug that inhibited 90% of the OD_580_ MTT colour conversion (IC_90_) compared to growth control wells with no drug added.

MBC was determined to discern bacteriostatic from bactericidal activities. Prior to MTT addition, 10 μL/well were transferred to 96-well LB agar plates and further incubated for 24 hours before addition of resazurin (30 μL/well). The MBC was defined as the lowest concentration of drug preventing a colour change from blue to pink. A compound was considered bactericidal if MBC/MIC ≤ 4 (17).

Solid MIC determinations were performed by the agar dilution method (18). Briefly, 24-well plates containing MHA with serial two-fold dilutions of tigecycline, colistin and fosfomycin were prepared in duplicates (V_F_= 1 mL/well). Upon agar solidification, each well was inoculated with 10 μL of a bacterial suspension (ca. 5×10^3^ CFU/well) and plates incubated for 18-20 hours. The MIC was considered as the lowest value that completely inhibited visible growth.

For fosfomycin susceptibility testing culture media was supplemented with 25 mg/L of glucose-6-phosphate according to EUCAST guidelines (20).

### Semi High-Throughput Synergy Screening (sHTSS)

The FDA library (n=1,430, 10 mM) was screened to identify synergistic partners of tigecycline, colistin and fosfomycin (Primary Compounds, PCs) against *K. pneumoniae* (17). Briefly, an overnight culture of *K. pneumoniae* was diluted to 10^5^ cells/mL in 22 mL of MHB medium containing 0.5% agar (top agar) and uniformly poured over 45 mL of MHB-1.5% agar (bottom agar) in OmniTrays (Nunc) in duplicates. PCs were added to the bottom agar at sub-inhibitory concentrations (MIC_sub_) of 0.125-0.25 mg/L (1/4x and 1/2xMIC) for tigecycline, 0.003-0.007 mg/L (1/256x and 1/128xMIC) for colistin and 16-32 mg/L (1/8x and 1/4xMIC) for fosfomycin. FDA-library compounds at 0.1-1 mM were transferred from 96-well tester plates onto top agar cell lawns using a pin replicator (1.6 mm pin diameter), which transferred approximately 200 nL/pin (0.2-2 nmol of each compound). OmniTrays were incubated overnight before inhibition zones measurement. Synergy was illustrated by an increase of the inhibition zones in the PC-containing agar plates (**Figure S1**).

Hits were classified in four categories: (i) synergy (Y): compounds whose inhibition zones increased at the two PCs MIC_sub_ tested (diameter MIC_sub1_ & diameter MIC_sub2_ > diameter_Control_); (ii) likely synergy (Y/N): compounds whose inhibition zones increased at only one PC MIC_sub_ (diameter MIC_sub1_ or diameter MIC_sub2_ > diameter_Control_); (iii) no interaction (N): compounds with no change in their inhibition zones (diameter MIC_sub1_ & diameter MIC_sub2_ = diameter_Control_); and, (iv) likely antagonism (A): compounds with decreased inhibition zones (diameter MIC_sub1_ & diameter MIC_sub2_ < diameter_Control_).

### Secondary validation assays

#### (i) Checkerboard assays

CBA were performed in 96-well plates using freshly prepared CAMHB. Each well was inoculated with 100 μL of 5×10^5^ CFU/mL (V_F_= 200 μL). Pre-inocula were prepared by direct suspension of bacteria grown overnight in MHB and plates incubated for 24 hours before determination of the compound activities alone and in combination (17,19). Fractional Inhibitory Concentration Indexes (FICI) were calculated as the sum of FIC_A_ plus FIC_B_ (FICI = FIC_A_ + FIC_B_) where FIC_A_ is the MIC of compound A in the presence of compound B divided by the MIC of compound A alone, and vice versa. Synergy was defined as FICI≤0.5, antagonism as FICI>4.0, and no interaction when the FICI was between 0.5-4.0 (21). Similarly, the Fractional Bactericidal Concentration Index (FBCI) was calculated as above described based on MBC values for each combination using the resazurin method (19).

#### (ii) Time-kill assays

Duplicates of exponentially growing *K. pneumoniae* cultures were inoculated in CAMHB 96-well plates (V_F_= 280 μL/well; 5×10^5^ CFU/mL) containing increasing compound concentrations (0.1x, 0.25x, 1x, 4x, 10x MIC values). At predefined time points (0, 2, 4, 6, 8, 24 and 48 hours), the bacterial population from each well was quantified by spot-platting 10-fold serial dilutions on MHA plates. Plates were incubated overnight and CFU/mL calculated. The lower limit of detection was 50 CFU/mL. Combo test concentrations were selected based on previous dose-response curves of the compound alone (typically 0.25x MIC and/or 1x MIC) or up to 64 mg/L in the case of inactive hits (**Figure S2**). MIC assays were run in parallel with the same inoculum as internal controls of compound activity. A synergistic combination was defined as a ≥2 log_10_ CFU/mL decrease in the bacterial count of the combination compared to the most active single agent at any time point (8, 24 and 48 hours). Antagonism was defined as a ≥2 log_10_ increase in CFU/mL between the combination and the most active single agent. All other cases were defined as indifferent. Bactericidal activity was defined as a ≥3 log_10_ CFU/mL reduction at any time point (8, 24 and 48 hours) compared to the initial inoculum (22).

## RESULTS

A general overview of the screening and validation progression cascade is displayed in **Figure 1. Table S2** displays a more comprehensive summary.

**Figure 1.**
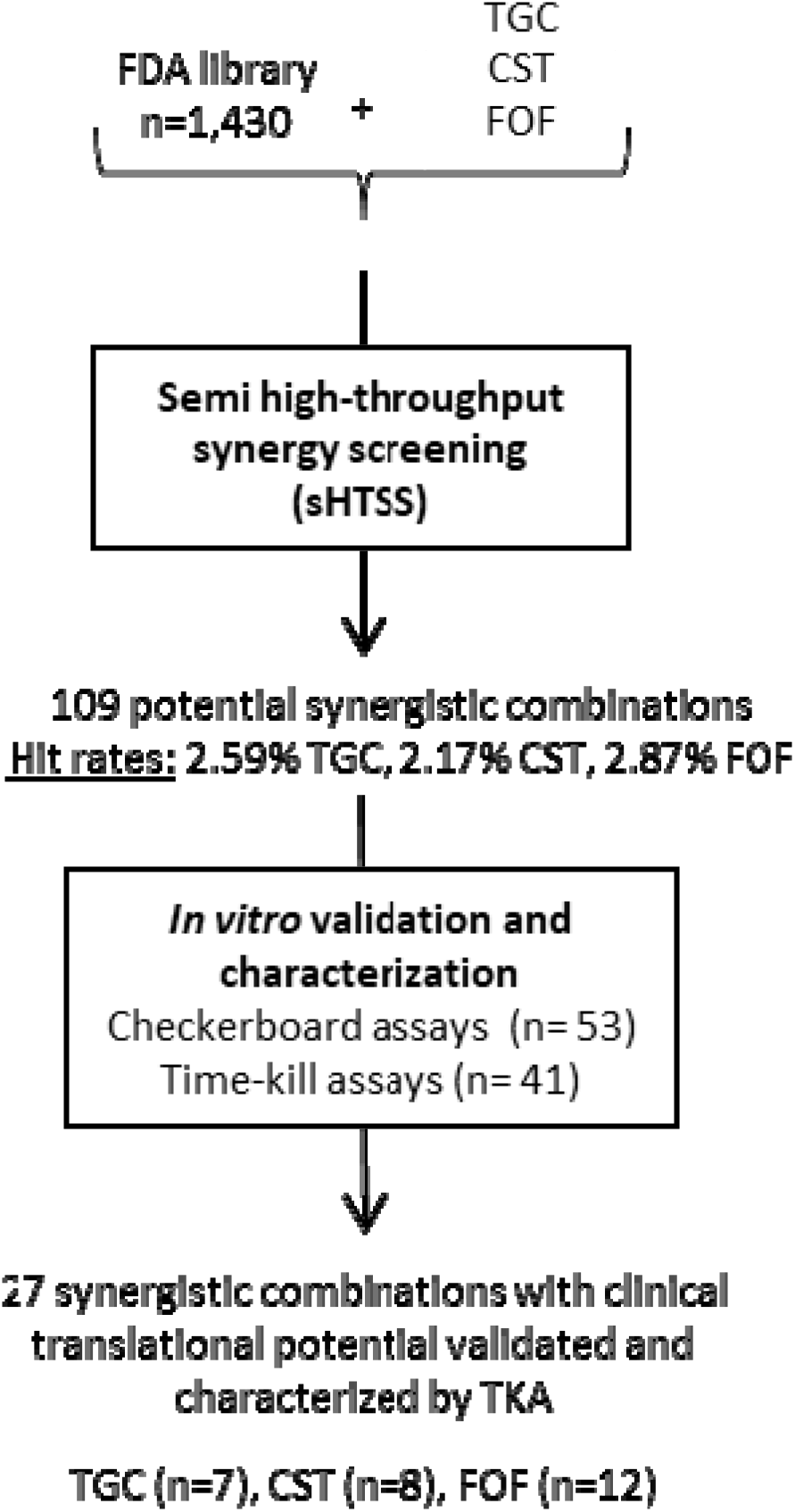
Progression cascade of the screening and validation activities performed in this study. The number of compounds tested, hit rates and hits validated are indicated at every step. TGC, tigecycline; CST, colistin; FOF, fosfomycin; TKA, time-kill assay.

### Synergy screenings of the FDA library in combination with last-line antibiotics against K. pneumoniae

We first determined the MIC of tigecycline, colistin and fosfomycin (0.5, 1 and 128 mg/L, respectively) on solid medium to select the two MIC_sub_ of each PC to be used in the three sHTSS (one per PC). Overall, 109 FDA compounds enhanced the activity of the PCs with hit discovery rates of 2.59% (n=37), 2.17% (n=31), and 2.87% (n=41) for tigecycline, colistin and fosfomycin, respectively. According to the interaction ranking criteria (see Materials & Methods) 19, 9 and 18 interactions were classified as synergistic, and 18, 22 and 23 as likely synergistic with tigecycline, colistin and fosfomycin, respectively. Additionally, 6, 10 and 12 interactions were classified as likely antagonistic with tigecycline, colistin and fosfomycin, respectively. We found some promiscuous compounds able to enhance the activity of more than one PC; thereby, among the 109 compounds that initially enhanced the activity of any of the PCs, there were 60 unique hits. A comprehensive hit list, inhibition zones and correlation with secondary validation assays are displayed in **Table S1**.

Classification by their therapeutic use revealed most were known antibiotics (75%), including quinolones (n=16), β-lactams (n=12) and aminoglycosides (n=6), among others. Non-antibiotic compounds (n=15) included other anti-infective agents (7%), antineoplastics (7%) or antipsychotics (3%) among others. These figures did not differ significantly among the three PCs (**Figure 2**).

**Figure 2.**
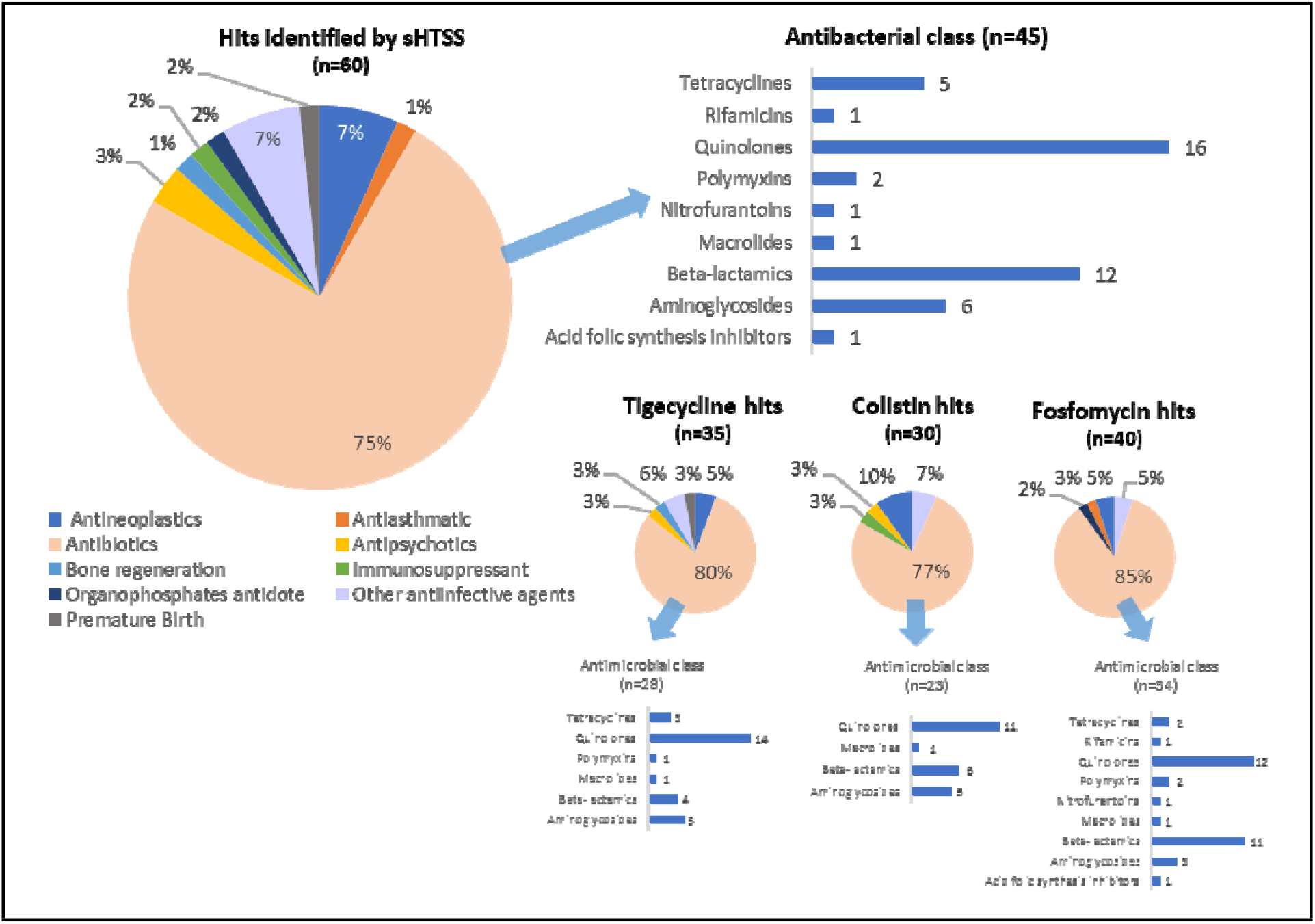
FDA compounds with favorable interactions with tigecycline, colistin and fosfomycin identified by sHTSS and classified by their therapeutic use. Other anti-infective agents include anti-parasitic, antiseptic and antiviral agents. Duplicate hits were removed from analysis. sHTSS, semi-high throughput synergy screening.

### Checkerboard assay displayed low validation rates

Fifty-three sHTSS interactions classified as synergy (n=50) or antagonism (n=3) for any PC were evaluated by CBA. FICI and FBCI indexes were calculated based on their MIC and MBC values respectively (**Table S3**); and interactions were validated in 8 out of the 53 (15.09%) combinations (**Table S1**). Colistin synergistic interactions were validated in 7 out of 12 combinations tested (58.33%) by both FICI and FBCI values. This percentage included the combination with bleomycin, classified as likely antagonism by sHTSS but showing synergy by CBA (FICI & FBCI = 0.16). Finally, synergy with fosfomycin was only validated in combination with lomefloxacin (1 out of 25, rate of 4%), with a FICI = 0.375 and FBCI = 0.365. No antagonism was confirmed by CBA (**Table S1**).

### Time-kill studies revealed novel promising combinations

Forty-one combinations (35 classified as Y or Y/N and 6 as A) were studied by TKA to assess their pharmacological and clinical translation potential (**Figure 3 & Table S4**). Overall, TKA showed a synergy validation rate of 65.85% (27 out of 41 combinations) at any of the predefined time points. Specific confirmation rates for each PC were 72.72% (8/11) for colistin, 70.58% (12/17) for fosfomycin, and 53.84% (7/13) for tigecycline.

**Figure 3.**
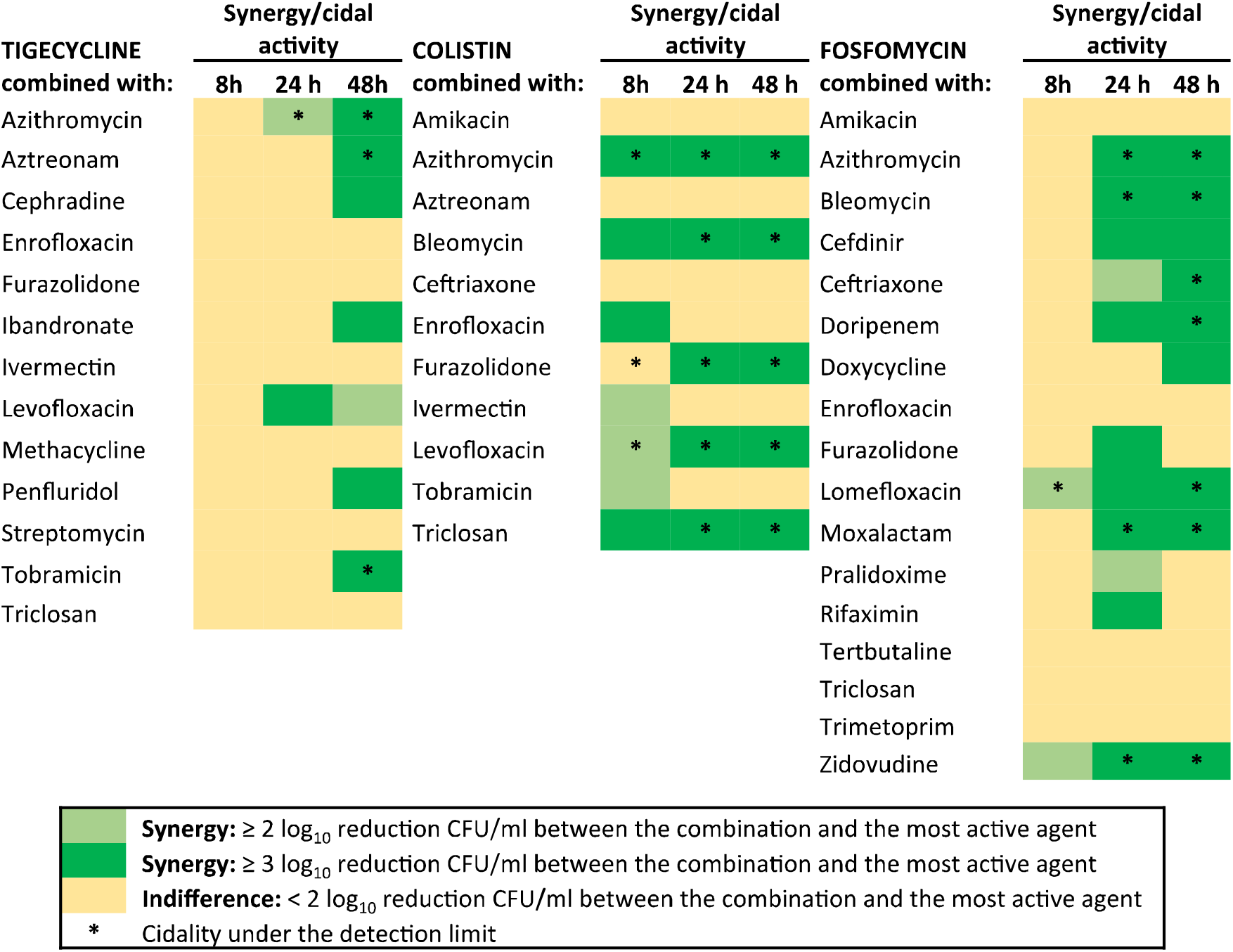
TKA drug interaction heat map. Forty-one sHTSS hits were prioritized for TKA based on a pharmacological and clinical translation potential assessment.

Six novel synergistic interactions with non-antibiotic drugs were confirmed in these studies (**Figure 4**). Strong bactericidal and synergistic interactions were observed between the antiviral zidovudine and fosfomycin, also in combinations of bleomycin (antineoplastic) with colistin and fosfomycin (**Figures 3 & 4**). Tigecycline combined with bisphosphonate ibandronate and the antipsychotic penfluridol showed a bacteriostatic profile and synergy at 48 hours with reductions of 4.04 log_10_ CFU/mL and 2 log_10_ CFU/mL, respectively. Both combinations prevented the bacterial regrowth observed with tigecycline alone (**Figure 4**). The antiparasitic ivermectin enhanced the activity of colistin at early time points (killing to the limit of detection at 5 hours) and showed synergistic effect at the 8-hour time point, although followed by bacterial rebound (**Figures 3 & 4**). Finally, pralidoxime (a poisoning antidote) showed synergy at 24 hours but no killing activity in combination with fosfomycin (**Table S4**).

**Figure 4.**
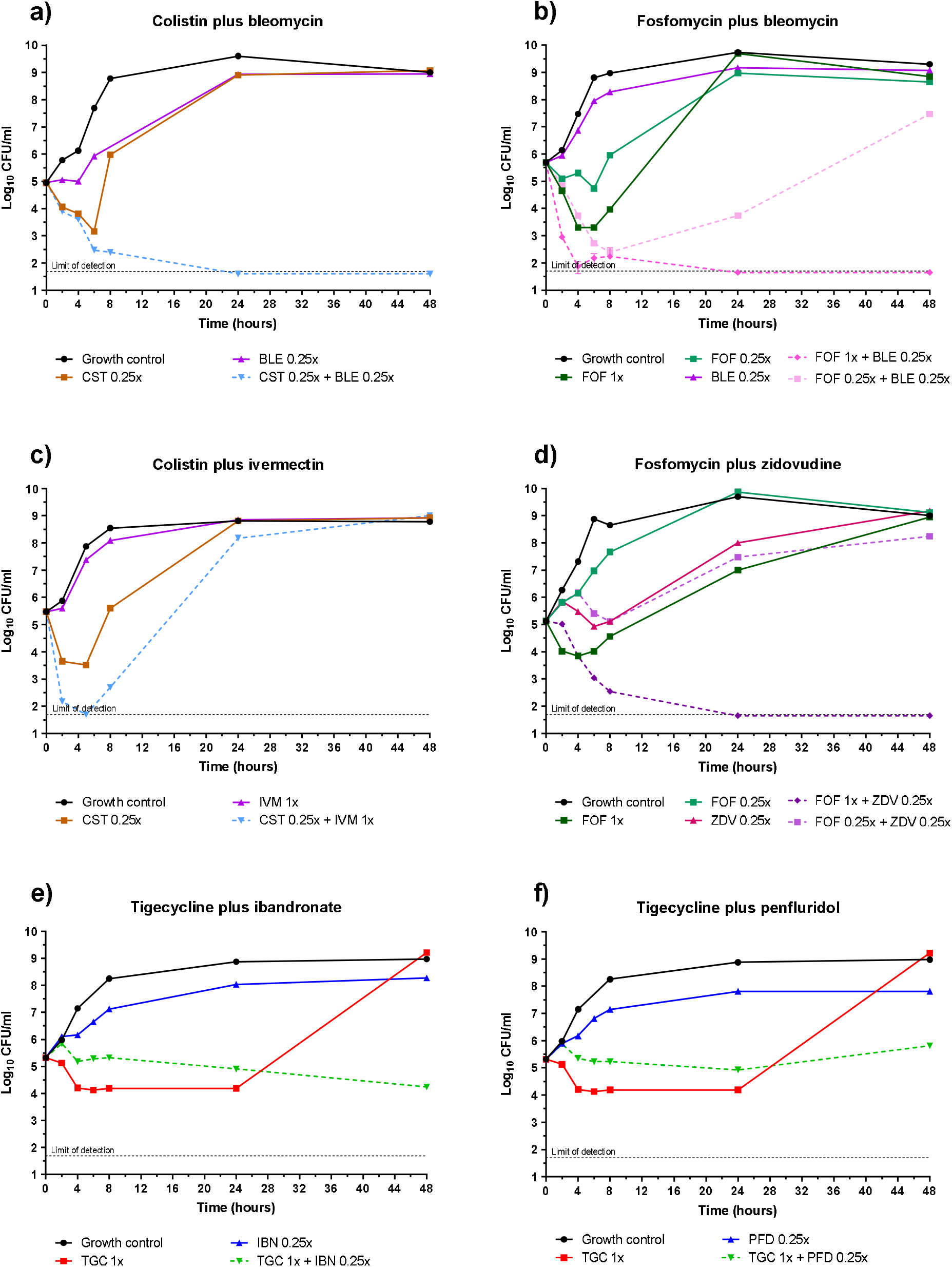
TKA of the novel combinations of non-antibiotic drugs with the PCs against *K. pneumoniae* ATCC 13883. **(a-b)** Bleomycin in combination with colistin or fosfomycin at subinhibitory concentration enhanced the bactericidal activity of both PCs **(c)** Ivermectin showed synergy up to 8 hours of incubation with colistin, after this time point a rebound was observed, similarly as both compounds alone **(d)** Zidovudine showed a strong interaction profile with fosfomycin at subinhibitory concentration (0.25x MIC) **(e-f)** Tigecycline combined with ibandronate or penfluridol showed static activity and synergy at 48 hours. BLE, bleomycin; CST, colistin; FOF, fosfomycin; IBN, ibandronate; IVM, ivermectin; PFD, penfluridol; TGC, tigecycline; ZDV, zidovudine.; MIC_BLE_ = 0.25 mg/L; MIC_CST_ = 1 mg/L; MIC_FOF_ ≥128 mg/L; MIC_IVM_ >64 mg/L (assumed in 64 mg/L); MIC_ZDV_ = 0.06 mg/L; MIC_TGC_ = 0.5 mg/L; MIC_IBN_ & MIC_PFD_ >32 mg/L (both assumed in 32 mg/L).

Effective combinations were also identified with other antimicrobials. Azithromycin displayed potent synergistic and bactericidal activities with all three PCs, showing the strongest effect in combination with colistin from the early 8-hour time point (**Figure 3**). A highly effective curve was also obtained with the colistin and levofloxacin combination. Moreover, we identified a remarkable potentiation of colistin by furazolidone and the antiseptic triclosan, which was not observed in the case of tigecycline or fosfomycin. Fosfomycin combinations with cefdinir, ceftriaxone, doripenem, lomefloxacin and moxalactam showed potent synergistic interactions at the 24- and 48-hour time points, the latter interaction reaching the limit of detection. Tigecycline showed bactericidal activity after 48 hours with aztreonam and tobramycin. Synergy with other antibiotics was also validated including cephradine, with a 2.35 log_10_ CFU/mL reduction at 48 hours; or levofloxacin, with synergy after 24 hours although regrowth was observed after 48 hours. An example of the type of information provided by the different validation assays (supporting the use of TKA over CBA) is shown in **Figure 5** for the tigecycline/aztreonam combination.

**Figure 5.**
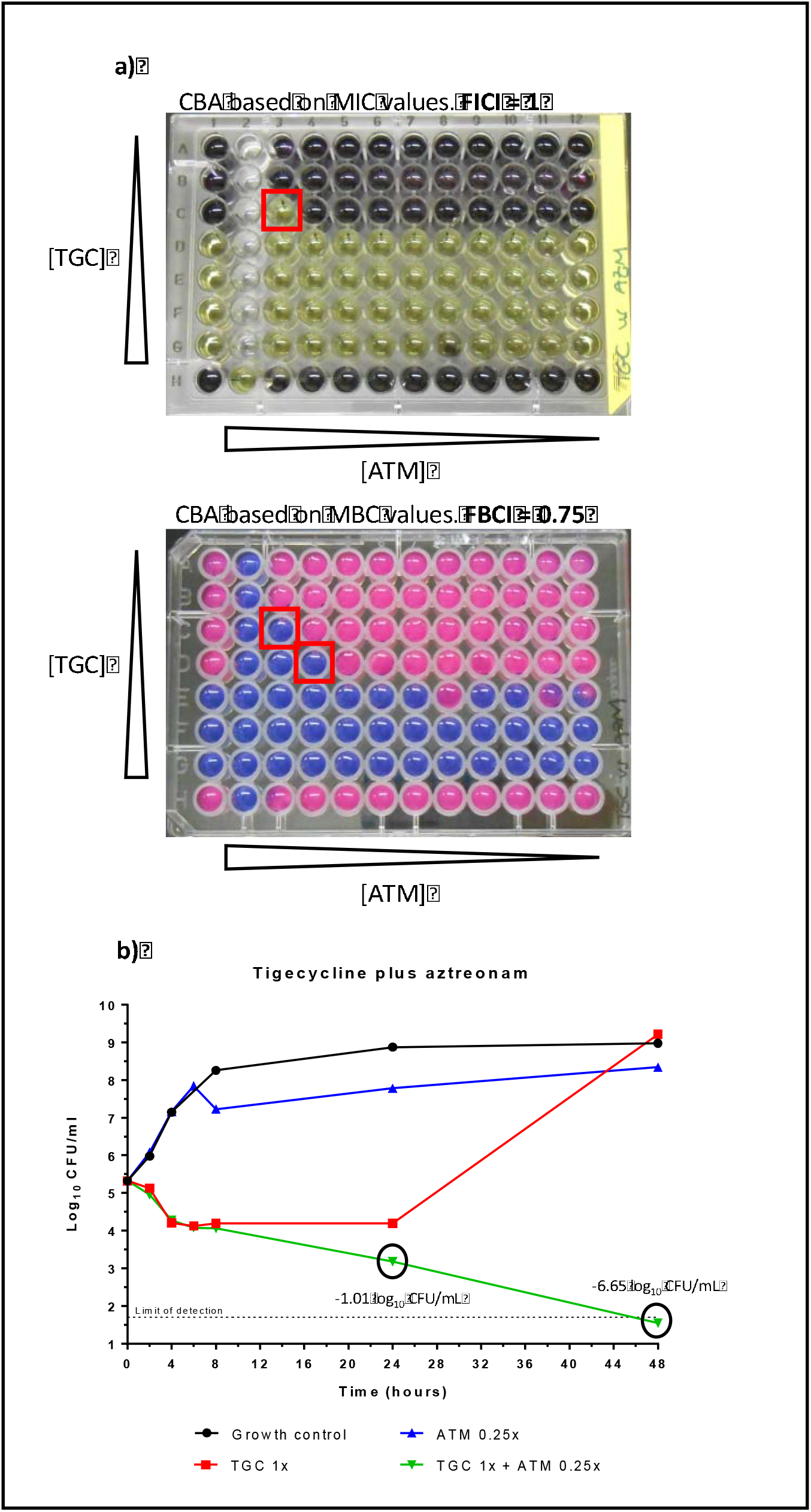
Secondary validation CBA and TKA of tigecycline in combination with aztreonam against *K. pneumoniae* ATCC 13883. **(a)** CBA were performed considering the MIC and MBC values of each compound alone and in combination. The FICI and FBCI were calculated from the most optimal combinatorial concentration (lowest value in the combination, red squares). Based on these assays, the combination was classified as “no interaction” (FICI/FBCI values between 0.5-4). **(b)** TKA provided longitudinal information showing a reduction of 1.01 log_10_ in CFU/mL of the combination with respect to tigecycline alone at 24 hours; thus, at this time point, the interaction was not classified as synergistic (<2 log_10_ reduction in CFU/mL with respect to the most active drug). However, after 48 hours the combination could be classified as synergistic with a reduction of 6.65 log_10_ CFU/mL with respect to tigecycline alone, and prevention of bacterial re-growth (proxy for sterilizing activity). MIC_ATM_ = 0.125-0.25 mg/L; MIC_TGC_ = 0.8 mg/L; MBC_ATM_ = 0.25 mg/L; MBC_TGC_ = 1.6 mg/L. CBA, checkerboard assay; TKA, time-kill assay; TGC, tigecycline; ATM, aztreonam.

## DISCUSSION

Traditional *in vitro* drug screening methodologies are tedious and time-consuming, and identification of novel combination therapies based on repurposed drugs could be a promising short/medium-term approach to fight against AMR. Synergy screenings (sHTSS) were initially developed to be used against mycobacteria and successfully identified novel combinations enhancing the activity of known antimicrobials (17,23). Here, we adapted the sHTSS methodology through a straightforward standardization process to find active combinations against *K. pneumoniae*. sHTSS demonstrated swiftly implementation for screening campaigns and allowed rapid, clear and simply drug interaction readouts derived from inhibition zones in agar media.

Our screenings in *K. pneumoniae* yield hit rates ranging from 2.17 to 2.87%, higher than that observed in *M. smegmatis* (1.4%) (17), but similar to other studies against Gram-negative bacilli with comparable hit rates (1.87%, tigecycline / 5.54%, colistin) (24). A large proportion of hits identified in our study were known antimicrobial drugs (82%), including antibiotics (75%) and other anti-infective agents (7%) (**Figure 2**). The overrepresentation of antimicrobials in synergy screening programs enhancing the activity of other antimicrobials is similarly described in other studies with *K. pneumoniae* (24,25) and *M. smegmatis* (17). Hind *et al.* observed this finding specially associated with *K. pneumoniae*, while they identified more heterogeneous targets with other Gram-negative bacteria (24).

Secondary validation assays performed by CBA and TKA provided discordant information; CBA yielded low sHTSS confirmation rates (15.09%), especially in the case of tigecycline and fosfomycin, while these increased to 65.85% when tested by TKA. Both techniques are widely used to determine synergy; however, they are based on different fundamental principles and parameters. Traditional CBA are based on growth inhibition parameters (MIC) at a fixed time point (usually overnight), while TKA reports bactericidal activity (Log_10_ CFU/mL) at several time points (up to 48 hours in our assays).

While performing CBA, we also measured the MBC (a fixed time point bactericidal parameter) to determine the FBCI, in addition to the FICI. This index has been largely disregarded in synergy studies, but some authors suggested FBCI might be a better predictor of drug interaction than FICI (26–28) and, in our view, it is a more stringent criteria to prioritize synergistic combinations. As an example, **Figure 5** shows validation results of the tigecycline/aztreonam combination. We found a clear “no interaction”profile by CBA (FICI=1), while a tendency towards synergism was observed by FBCI, which some authors interpreted as additivity (0.5 < FBCI ≤ 1), although this terminology should be avoided (21). The use of a fixed time point bactericidal parameter could predict TKA data; the combination showed no synergy at 24 hours (although with a tendency to reduce the bacterial count compared to tigecycline alone) but prevented growth rebound at 48 hours, suggesting a synergistic interaction. Nevertheless, the use of the FBCI might come with some limitations for compounds with high MIC values (≥128 mg/L) such as fosfomycin, for which we could not calculate the FBCI because the MBC exceeded the maximum tested concentration.

The fixed time point limitation of the MBC assays is addressed in TKA by the inclusion of longitudinal pharmacodynamic data, which allows TKA as a most suitable methodology to robustly characterize drug interaction dynamics, a paradigm shift in antimicrobial development methodologies (29). We decided to perform extended TKA up to 48 hours of incubation with the aim to prioritize the most effective combinations that maintained bactericidal activities up to the end of the assay. Moreover, the 48-hour time point allow us to identify combinations that could potentially lead to therapeutic failure when used in the clinical practice, i.e., combinations considered initially effective but that rebounded after the 24-hour time point (**Figure S2**). We believe this could be a better proxy of sterilization and, thus, potency of the combination.

Out of the three PCs used in this study, the highest number of validated combinations was obtained for colistin and fosfomycin, antibiotics targeting the outer membrane and cell wall, respectively. Both antibiotics have been associated with enhance killing when in combination with intracellular targeting compounds, explained by an increase in permeability (24,30,31). In agreement with our study, clinically relevant synergistic interactions with our three PCs have been reported within a combinatorial therapy for the treatment of MDR enterobacteria (32,33). Moreover, synergy was demonstrated between tigecycline and aminoglycosides (34–36), colistin and levofloxacin (37,38), fosfomycin and cephalosporins (31) or doripenem (39,40) against *K. pneumoniae*. Principe *et al.* also reported synergy between tigecycline and levofloxacin against *Acinetobacter baumanii* strains (41). A recent review on *in vitro* fosfomycin combinations supports our findings in which we did not observe synergy between fosfomycin and amikacin or trimethoprim (42). Using CBA, Ontong *et al.* reported synergy between colistin/amikacin and colistin/tobramycin for 72.72% and 45.45% of MDR *K. pneumoniae* strains, respectively (37). We also observed synergy by CBA for the latter combination, but such interactions were not maintained by TKA (**Tables S1 & S4**). This highlights again TKA as a much better proxy than CBA to identify synergistic combinations.

Novel combinations including non-antibiotics were identified: (i) *Tigecycline plus ibandronate or penfluridol*. Ibandronate and other bisphosphonate derivatives are anti-parasitic drugs inhibiting the synthesis of essential isoprenoids (43,44). The antipsychotic penfluridol displayed partial synergistic activity with aminoglycosides and β-lactams against *Enterococcus faecalis* (45). To the best of our knowledge, here we report for the first time the antibacterial activity of both drugs with tigecycline against enterobacteria; (ii) *Ivermectin with tigecycline or colistin.*Ivermectin is an anthelmintic with both veterinary and clinical applications, widely explored for different repurpose anti-parasitic and antiviral uses (46–48). Its antimicrobial activity was demonstrated against *Mycobacterium* species (49–51). sHTSS classified ivermectin as likely antagonism with tigecycline and colistin. Follow up TKA validation studies demonstrated no interaction with tigecycline and a slight synergistic effect at the 8-hour time point in combination with colistin but followed by a bacterial rebound, highlighting the need to perform secondary CFU-based validation assays (**Figures 3 & 4**). (iii) *Zidovudine plus fosfomycin*. Zidovudine was the first antiretroviral used for HIV treatment. Since 1980s, its antibacterial activity against enterobacteria is attributed to its targeting of bacterial thymidine kinases (52). Several *in vitro* and/or *in vivo* studies demonstrated effective antibacterial activities of zidovudine in combination with antibiotics such as fluoroquinolones, trimethoprim, aminoglycosides, tigecycline and polymyxins (52–57). Similar to a recent report (58), we also identified a strong synergistic interaction with fosfomycin; (iv) *Bleomycin plus colistin or fosfomycin*. The cytotoxicity of bleomycin poses a barrier to an antibacterial repurposing approach; nevertheless, our results could set up the basis for the development of analogues or dosage-based studies to minimize toxicity; (v) *Azithromycin*. We validated the synergistic interaction of azithromycin in combination with all three PCs. This was in agreement with several studies reporting the bactericidal and antibiofilm action of azithromycin combined with tigecycline (59), and colistin (60,61) against different Gram-negative bacilli. The interaction with fosfomycin is novel. Another macrolide, erythromycin, displayed synergy for 50% of the *K. pneumoniae* strains tested (62). The promiscuity of azithromycin interactions suggests its inclusion as primary compound in future synergy screening campaigns, which might ease the identification of new synergistic partners to improve its therapeutic use.

In summary, we have identified novel synergistic combinations against *K. pneumoniae* adapting the sHTSS screening methodology and using novel validation endpoints in TKA. In this work, we have evaluated the efficacy of the interactions against a *K. pneumoniae* reference strain with high susceptibility to most antibiotics, which might overestimate the real rate of effective combinations against clinical and MDR strains. We expect this scenario could be unlikely, and this technology should be helpful as well for readily identifying synergistic combinations active against MDR strains. Moreover, both CBA or TKA are methodologies in which drugs are added only once at the start of the experiment at fixed concentrations; although this is a technical simplification of the dynamic fluctuation of drugs in the clinical practice, other *in vitro* methodologies such as the hollow fibre system computing the pharmacokinetic properties of the compounds in combination and linking it to pharmacodynamics parameters of activity might provide essential information to refine drug doses and hence aid the design of future clinical trials.

Finally, clinical implementation of synergistic novel combinations might improve medical decision-making; the combined effect could reduce the exposure to potentially toxic drugs, resulting in lower incidence of treatment-related side-effects and complications, but enhancing their effectiveness.

## Acknowledgements

We thank Ana Isabel López-Calleja and Antonio Rezusta from the “Servicio de Microbiología, Hospital Universitario Miguel Servet, Zaragoza, Spain” for helpful discussions during the performance of this work and critical reading of the manuscript.

## Data availability statement

All data pertaining to this work is within the main manuscript or supplementary information. Primary data are available from the corresponding author upon request.

