## Supplementary Data for "Novel synergistic combinations of last-line antibiotics and FDA-approved drugs against Klebsiella pneumoniae revealed by in vitro synergy screenings"

Gómara-Lomero, M. *et al.*

**SUPPLEMENTARY FIGURES**

**Figure S1. Representative plate analysis of a semi-High-Throughput Synergy Screen (sHTSS).** FDA compounds were pin-spotted onto a soft agar lawn of *K. pneumoniae* in the absence (control plate) or in the presence of subinhibitory concentrations of fosfomycin (16 and 32 mg/L; 1/8xMIC and 1/4xMIC, respectively). Compounds whose zones of inhibition were larger in the presence of fosfomycin than in plates without fosfomycin (examples A, B or C) were selected as hits for further validation of their potential synergistic interaction. High-density plates overlapping inhibition zones for two or more compounds (D, inhibition zones >15 mm) were deconvoluted at a lower compound density or lower concentration (0.1 mM) for clear inhibition zone readings. Ø, inhibition zone diameter value; Red arrows, inhibition zone diameter; r, inhibition zone radius (when diameter cannot be determined); MIC_FOF_= 128 mg/L.

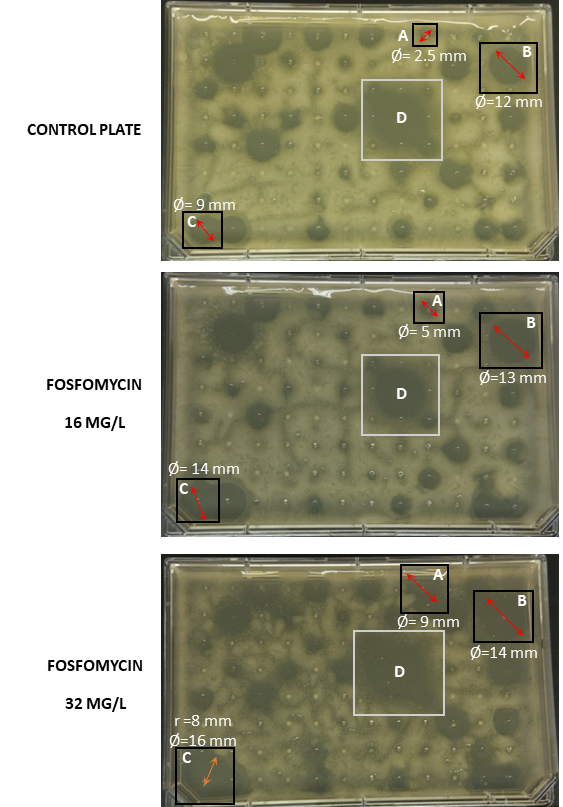

**Figure S2. Representative time-kill assays of drugs alone and in combination against *K. pneumoniae* ATCC 13883. (a-f)** Dose response curves (0.1x, 0.25x, 1x, 4x and 10xMIC) of compounds alone; **(g-i)** Combination time-kill curves were performed at pairing static and/or sub-inhibitory concentrations (0.25x and/or 1xMIC values); **(g)** Synergistic interaction with killing activity up to 24 hours followed by a bacterial growth rebound; **(h)** No interaction pattern; **(i)** Synergistic interaction with bactericidal activity at both 24 and 48 hours. Synergy was interpreted as a ≥2 log_10_ CFU/mL decrease in bacterial count compared to the most active single agent in the combination at 8, 24 and 48 hours. Bactericidal activity was interpreted as a ≥3 log_10_ CFU/mL reduction at 8, 24 and 48 hours compared to the initial inoculum. An untreated growth control was always included. MIC_TGC_= 0.5 mg/L; MIC_LVX_= 0.03 mg/L; MIC_CST_= 1 mg/L; MIC_AMK_= 1 mg/L; MIC_FOF_= 128 mg/L; MIC_DOR_= 0.03 mg/L. TGC, tigecycline; LVX, levofloxacin; CST, colistin; AMK, amikacin; FOF, fosfomycin; DOR, doripenem.

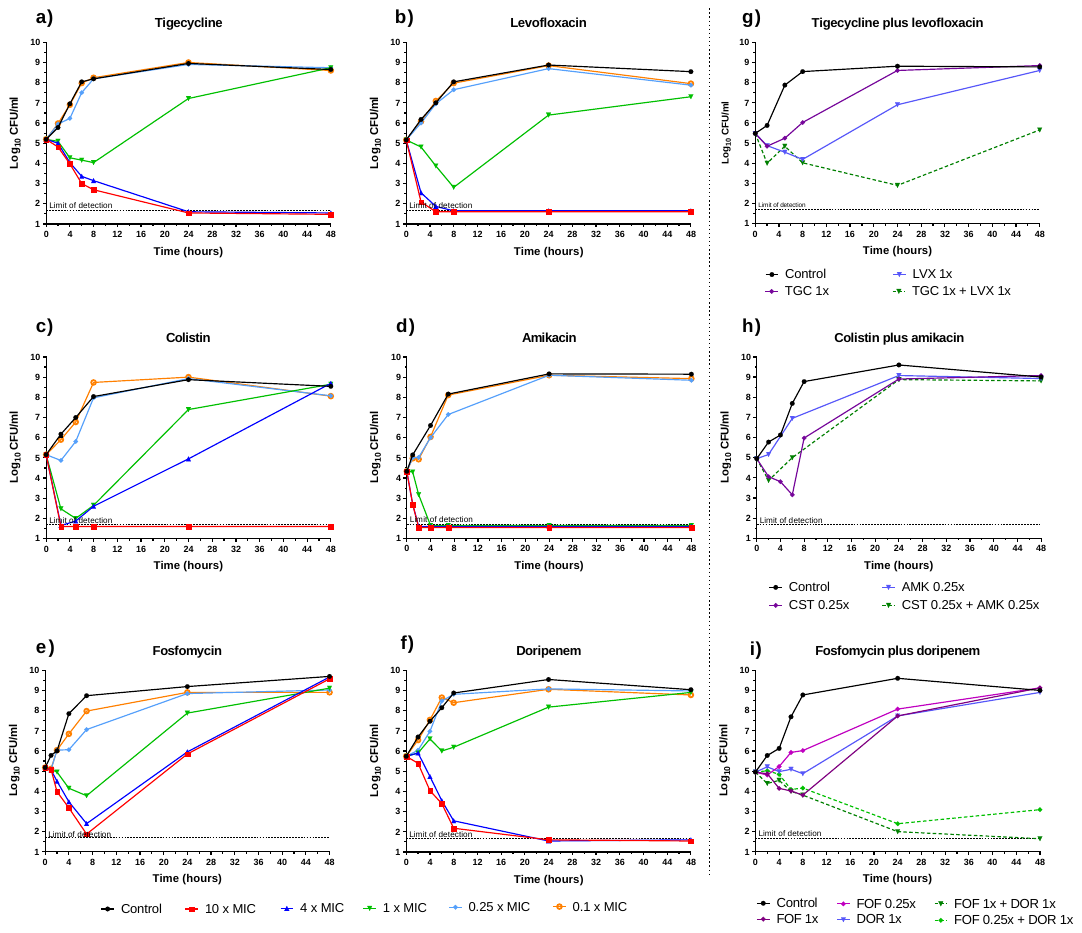

**SUPPLEMENTARY TABLES**

**Table S1. FDA compounds identified in sHTSS with tigecycline, colistin and fosfomycin and validation data against *K. pneumoniae* ATCC 13883**

See supplementary Table S1 file.

**Table S2. Results obtained in the screening and validation of synergistic combinations against K. pneumoniae ATCC 13883.**

|  | **Primary Compound** | **Tigecycline** | | | **Colistin** | | | **Fosfomycin** | | | **Grand total (three PCs)** | | |
| --- | --- | --- | --- | --- | --- | --- | --- | --- | --- | --- | --- | --- | --- |
|  |  | **S** | **A** | **Total** | **S** | **A** | **Total** | **S** | **A** | **Total** | **S** | **A** | **Total** |
|  | **sHTSS^a^** | 37^*^ | 6 | 43 | 31^*^ | 10 | 41 | 41^*^ | 12 | 53 | **109** | **28** | 137 |
| **Secondary** **validation** | **CBA assayed** | 14 | 2 | 16 | 11 | 1 | 12 | 25 | 0 | 25 | 50 | 3 | 53 |
|  | **CBA validated^b^** | 0 | 0 | 0 | 6 + 1^#^ | - | 7 | 1 | 0 | 1 | 8 | 0 | 8 |
|  | **TKA assayed** | 10 | 3 | 13 | 8 | 3 | 11 | 17 | 0 | 17 | 35 | 6 | 41 |
|  | **TKA validated^c^** | 7 | 0 | 7 | 5 + 3^#^ | - | 8 | 12 | 0 | 12 | 27 | 0 | 27 |
| **Novelty** | **Already published combinations** | 3 | - | - | 3 | - | - | 2 | - | **-** | 8 | - | - |
|  | **Novel combinations (non-antibiotics)** | 2 | - | - | 2 | - | - | 3 | - | **-** | 7 | - | - |
|  | **Novel combinations (other)** | 2 | - | - | 3 | - | - | 7 | - | **-** | 12 | - | - |

**^a^**Interaction classification was based on the increment of the inhibition zones at the two PCs sub-inhibitory concentrations tested compared to those of the no PC plates as described in Material and Methods. Raw data are displayed, including compounds with different chemical forms (i.e., amikacin hydrate / amikacin disulfate)

^b^FICI or FBCI ≤0.5 indicates synergy

^c^Synergy was defined as a ≥2 log_10_ reduction in CFU/mL in the combination compared to the most active compound alone

^*^Synergistic interactions included those classified as synergy (Y) and likely synergy (Y/N)

^#^Interaction validated as synergy by CBA or TKA although sHTSS initially identified antagonistic interaction

S: synergy; A; antagonism; sHTSS, semi-high throughput synergy screening; CBA, checkerboard assays; TKA, time-kill assays

**Table S3. Drug susceptibility of *K. pneumoniae* ATCC 13883 to FDA compounds presenting synergy with tigecycline, colistin or fosfomycin*.*** ^a^Determined by MTT assay. ^b^Determined by resazurin assay.

| **Compound** | **MIC values (mg/L)^a^** | **MBC values (mg/L)^b^** |
| --- | --- | --- |
| Amikacin | 1 | 2 |
| Azithromycin | 2 | 2-4 |
| Aztreonam | 0.125 | 0.25 |
| Balofloxacin | 0.25 | 0.25 |
| Bleomycin | 0.25 | 0.25 |
| Cefdinir | 0.5 | 0.5 |
| Cefmenoxime | 0.25 | 0.25-0.5 |
| Cefoperazone | 4 | 4 |
| Ceftazidime | 0.5 | 0.5 |
| Ceftiofur | 1 | 1 |
| Ceftriaxone | 0.125 | 0.125 |
| Cephradine | 32 | 32 |
| Colistin | 1-2 | 1-2 |
| Danofloxacin | 0.06 | 0.12 |
| Difloxacin | 0.12-0.25 | 0.25-0.5 |
| Doripenem | 0.03 | 0.06 |
| Doxycycline | 4 | >8 |
| Enrofloxacin | 0.015-0.03 | 0.03-0.06 |
| Flumequine | 1-2 | 1-2 |
| Fosfomycin | ≥128 | ≥128 |
| Furazolidone | 1 | 1 |
| Ibandronate | >32 | >32 |
| Ivermectin | >64 | >64 |
| Levofloxacin | 0.03 | 0.03 |
| Lomefloxacin | 0.12-0.25 | 0.12 |
| Marbofloxacin | 0.015-0.03 | 0.015-0.06 |
| Methacycline | 1 | 32 |
| Moxalactam | 0.5 | 0.5 |
| Moxifloxacin | 0.25-0.5 | 0.25-0.5 |
| Nadifloxacin | 1 | 1 |
| Netilmicin | 1 | 1 |
| Norfloxacin | 2 | 2 |
| Ofloxacin | 0.125 | 0.125 |
| Pefloxacin | 0.125 | 0.125 |
| Penfluridol | >32 | >32 |
| Pralidoxime | >32 | >32 |
| Rifaximin | 16 | 32 |
| Sisomicin | 0.125 | 0.25 |
| Sparfloxacin | 0.03 | 0.03 |
| Streptomycin | 2 | 2 |
| Terbutaline | >32 | >32 |
| Tigecycline | 0.5 | 1? |
| Tobramicin | 0.125 | 0.125 |
| Triclosan | 0.25-0.5 | 0.25 |
| Trimetoprim | 1 | 8-16 |
| Zidovudine | 0.06 | 0.5-0.12 |

**Table S4. Time-kill assays drug interaction data against *K. pneumoniae* ATCC 13883**. This table includes the raw data supporting Figure 3. Data display the number of residual viable colonies (Δlog_10_ CFU/mL) between the combination and the most active agent alone and between the initial and final inoculum at the different time points (after 8, 24 and 48 hours of incubation). A negative sign denotes a reduction in the viable counts. Values in bold: synergistic (≥2 log_10_ CFU/mL reduction) and bactericidal effects (≥3 log_10_ CFU/mL reduction). Initial inoculum was 5x10^5^ CFU/mL. The experimental limit of detection was 50 CFU/mL. AMK, amikacin; ATM, aztreonam; AZM, azithromycin, BLE, bleomycin; BUT, tertbutaline; CDR, cefdinir; CRO, ceftriaxone; CST, colistin; DOX, doxycycline; DOR, doripenem; ENR, enrofloxacin; FOF, fosfomycin; FZD, furazolidone; IBN, ibandronate; IVM, ivermectin; LVX, levofloxacin; LMF, lomefloxacin; MET, methacycline; MOX, moxalactam; PFD, penfluridol; PRA, pralidoxime; RAD, cephradine; RFX, rifaximin; STP, streptomycin; TCS, triclosan; TGC, tigecycline; TMP, trimethoprim; TOB, tobramycin; ZDV, zidovudine.

|  |  | **Δlog_10_ CFU/mL between the combination and the most active agent** | | |  | **Δlog_10_ CFU/mL between initial and final inoculum** | | |
| --- | --- | --- | --- | --- | --- | --- | --- | --- |
| **Combinations** | **Concentrations (mg/L)** | **8 h** | **24 h** | **48 h** |  | **8 h** | **24 h** | **48 h** |
| **TGC / AZM** | 0.5 / 2 | -0.99 | **-2.50** | **-6.01** |  | -2.12 | **-3.63** | **-3.63** |
| **TGC / ATM** | 0.5 / 0.025 | -0.13 | -1.01 | **-6.65** |  | -1.26 | -2.15 | **-3.63** |
| **TGC / RAD** | 0.5 / 8 | -0.08 | -1.24 | **-5.93** |  | -1.21 | -2.37 | -2.34 |
| **TGC / ENR** | 0.5 / 0.003 | 0.36 | -0.69 | -0.74 |  | -1.35 | 0.93 | 2.26 |
| **TGC / FZD** | 0.5 / 0.25 | 2.48 | -0.70 | 0.03 |  | -0.82 | 2.43 | 3.40 |
| **TGC / IBN** | 0.5 / 8 | 1.13 | 0.71 | **-4.04** |  | 0 | -0.42 | -1.08 |
| **TGC / IVM** | 0.5 / 64 | -0.07 | -0.44 | 0 |  | 0.48 | 2.68 | 3.37 |
| **TGC / LVX** | 0.5 / 0.03 | -0.17 | **-4.00** | **-2.95** |  | -1.46 | -2.57 | 0.18 |
| **TGC / MET** | 0.5 / 1 | -0.04 | -0.32 | -1.74 |  | -1.18 | -1.45 | 2.15 |
| **TGC / PFD** | 0.5 / 8 | 1.04 | 0.74 | **-2.00** |  | -0.09 | -0.39 | 0.49 |
| **TGC / STP** | 0.5 / 0.5 | -0.91 | -0.47 | -0.73 |  | -0.36 | 2.65 | 2.40 |
| **TGC / TOB** | 0.5 / 0.125 | -0.09 | -1.19 | **-7.23** |  | -1.23 | -2.32 | **-3.63** |
| **TGC / TCS** | 0.5 / 0.125 | -1.94 | -0.43 | -0.03 |  | -1.40 | 2.70 | 3.34 |
| **CST / AMK** | 0.25 / 0.125 | -1.98 | -0.03 | -0.12 |  | -0.95 | 3.92 | 3.86 |
| **CST / AZM** | 0.25 / 2 | **-4.29** | **-6.29** | **-5.24** |  | **-3.26** | **-3.26** | **-3.26** |
| **CST / ATM** | 0.25 / 0.025 | 0.02 | -0.04 | 0.12 |  | 1.05 | 3.07 | 3.65 |
| **CST / BLE** | 0.25 / 0.06 | **-3.58** | **-7.21** | **-7.26** |  | -2.56 | **-3.26** | **-3.26** |
| **CST / CRO** | 0.25 / 0.03 | 0.14 | 0.10 | -0.10 |  | 1.16 | 2.89 | 2.95 |
| **CST / ENR** | 0.25 / 0.003 | **-3.98** | -1.45 | -0.38 |  | -2.95 | 1.44 | 3.21 |
| **CST / FZD** | 0.25 / 0.25 | -0.49 | **-7.09** | **-7.24** |  | **-3.79** | **-3.79** | **-3.79** |
| **CST / IVM** | 0.25 / 64 | **-2.91** | -0.64 | 0.07 |  | -2.78 | 2.70 | 3.52 |
| **CST / LVX** | 0.25 / 0.03 | **-2.50** | **-5.21** | **-6.92** |  | **-3.79** | **-3.79** | **-3.79** |
| **CST / TOB** | 0.25 / 0.125 | **-2.65** | -0.72 | -0.56 |  | -2.52 | 2.62 | 2.43 |
| **CST / TCS** | 0.25 / 0.125 | **-3.68** | **-7.21** | **-7.39** |  | -2.65 | **-3.26** | **-3.26** |
| **FOF / AMK** | 128 / 0.125 | -0.36 | -0.84 | 0.20 |  | -2.10 | 2.16 | 3.34 |
|  | 32 / 0.125 | -0.03 | -1.05 | 1.05 |  | 0.23 | 2.23 | 4.00 |
| **FOF / AZM** | 128 / 2 | -1.01 | **-6.18** | **-4.37** |  | -2.74 | **-4.01** | **-4.01** |
|  | 32 / 2 | -1.49 | **-6.18** | **-4.37** |  | -1.22 | **-4.01** | **-4.01** |
| **FOF / BLE** | 128 / 0.06 | -1.79 | **-7.01** | **-7.15** |  | **-3.52** | **-4.01** | **-4.01** |
|  | 32 / 0.06 | **-3.57** | **-5.24** | -1.18 |  | **-3.30** | -1.96 | 1.78 |
| **FOF / CDR** | 128 / 0.125 | -1.82 | **-5.31** | **-6.75** |  | -2.39 | **-3.44** | -2.95 |
|  | 32 / 0.125 | -0.87 | -0.22 | -0.23 |  | -1.20 | 2.52 | 3.57 |
| **FOF / CRO** | 128 / 0.03 | -0.54 | **-2.12** | **-6.24** |  | -1.11 | -0.26 | **-3.44** |
|  | 32 / 0.03 | **-2.61** | 0.37 | 0.63 |  | -0.07 | 2.71 | 3.43 |
| **FOF / DOR** | 128 / 0.03 | 0 | **-5.74** | **-7.21** |  | -1.14 | -2.95 | **-3.26** |
|  | 32 / 0.03 | -0.71 | **-5.34** | **-5.81** |  | -0.79 | -2.56 | -1.86 |
| **FOF / DOX** | 128 / 4 | 1.03 | -1.71 | **-3.85** |  | -0.11 | -2.18 | -0.48 |
|  | 32 / 4 | 0.05 | -0.94 | -0.40 |  | 0 | -1.41 | 2.98 |
|  | 128 / 1 | 0.27 | **-2.72** | -0.43 |  | -0.88 | 0.07 | 3.74 |
|  | 32 / 1 | -0.12 | -0.10 | 0.03 |  | 0.95 | 3.02 | 4.21 |
| **FOF / ENR** | 128 / 0.003 | -1.61 | -1.07 | 0.11 |  | -2.18 | 0.80 | 3.93 |
|  | 32 / 0.003 | -1.61 | -0.20 | 0.15 |  | 0.93 | 3.32 | 4.06 |
| **FOF / FZD** | 128 / 0.25 | -1.56 | **-4.52** | -1.11 |  | -2.13 | -2.65 | 2.71 |
|  | 32 / 0.25 | 0.30 | 0 | -0.40 |  | 1.02 | 3.17 | 3.57 |
| **FOF / LMF** | 128 / 0.025 | **-2.27** | **-6.07** | **-7.15** |  | **-4.01** | **-3.22** | **-4.01** |
|  | 32 / 0.025 | -0.79 | -1.57 | 0.28 |  | -0.52 | 1.28 | 3.23 |
| **FOF / MOX** | 128 / 0.125 | -0.27 | **-4.62** | **-6.62** |  | -1.41 | **-3.26** | **-3.26** |
|  | 32 / 0.125 | 0.19 | 1.17 | 0.8 |  | -0.86 | 2.52 | 4.16 |
| **FOF / PRA** | 128 / 8 | -0.23 | -0.10 | 0.30 |  | -0.80 | 1.77 | 4.12 |
|  | 32 / 8 | -0.55 | **-2.46** | 0.43 |  | 1.98 | 1.89 | 4.41 |
| **FOF / RFX** | 128 / 4 | 0.12 | **-4.90** | -1.80 |  | -1.02 | -2.11 | 2.26 |
|  | 32 / 4 | -0.24 | -0.12 | -0.05 |  | 0.82 | 3.00 | 4.12 |
| **FOF / BUT** | 128 / 8 | 0.39 | 1.00 | 0.24 |  | -0.18 | 2.87 | 4.06 |
|  | 32 / 8 | 0.21 | -0.15 | 0.02 |  | 2.74 | 3.85 | 4.00 |
| **FOF / TCS** | 128 / 0.125 | 0.64 | 1.18 | -0.30 |  | 0.07 | 3.05 | 3.52 |
|  | 32 / 0.125 | -0.51 | -0.30 | 0.49 |  | 1.89 | 3.47 | 4.47 |
| **FOF / TMP** | 128 / 0.5 | -0.52 | -0.43 | -1.93 |  | -1.09 | 1.44 | 1.89 |
|  | 32 / 0.5 | -0.55 | -1.22 | -1.00 |  | 1.98 | 2.39 | 2.89 |
| **FOF / ZDV** | 128 / 0.015 | **-2.02** | **-5.31** | **-7.26** |  | -2.59 | **-3.44** | **-3.44** |
|  | 32 / 0.015 | 0 | -0.53 | -0.87 |  | -0.02 | 2.35 | 3.11 |
