## Supplementary Table S1 for "Novel synergistic combinations of last-line antibiotics and FDA-approved drugs against Klebsiella pneumoniae revealed by in vitro synergy screenings"

**Table S1.** **FDA compounds identified in sHTSS with tigecycline, colistin and fosfomycin and validation data against K. pneumoniae ATCC 13883**

| ***TIGECYCLINE HITS*** | | **sHTSS** | | | |  |  |  |  |  |  |  |  |
| --- | --- | --- | --- | --- | --- | --- | --- | --- | --- | --- | --- | --- | --- |
|  |  |  | **Δ zones of inhibition (mm) ^b^** | |  | **CBA^e^** | | **TKA^f^** | | |  |  |  |
| **PubChem #^a^** | **FDA compound** | **Ø^c^** | **TGC ¼ x MIC** | **TGC ½ x MIC** | **Interaction^d^** | **FICI** | **FBCI** | **8h** | **24h** | **48h** | **Chemical classification^g^** | **Therapeutic use** | **Target** |
| CID 441384 | Abacavir sulfate | 0 | 0 | 1.6 | Y/N | - | - | - | - | - | Nucleoside analogue | Antiretroviral | Reverse transcriptase |
| CID 38351 | Amikacin disulfate | 0 | 0 | 3 | Y/N | 0.5 | 0.75 | - | - | - | Aminoglycoside | Antibiotic | 30S ribosomal |
| CID 16218899 | Amikacin hydrate | 0 | 0 | 2.65 | Y/N | 0.5 | 0.75 | - | - | - | Aminoglycoside | Antibiotic | 30S ribosomal |
| CID 87665603 | Atosiban Acetate | 0 | 0 | 1.65 | Y/N | - | - | - | - | - | Oligopeptide | Premature Birth | Oxytocin receptor |
| CID 447043 | Azithromycin | 0 | 2.5 | 6 | Y | 0.5-4 | 0.5-4 |  | * | * | Macrolide | Antibiotic | 50S ribosomal |
| CID 3033819 | Azithromycin Dihydrate | 4.7 | 1.45 | 1.75 | Y | 0.5-4 | 0.5-4 |  | * | * | Macrolide | Antibiotic | 50S ribosomal |
| CID 5742832 | Aztreonam | 11 | 3 | 4 | Y | 1 | 0.75 |  |  | * | Monobactam | Antibiotic | PBP 3 |
| CID 65958 | Balofloxacin | 3.5 | 1 | 0 | Y/N | - | - | - | - | - | Fluoroquinolone | Antibiotic | DNA gyrase |
| CID 72466 | Bleomycin Sulfate | 9.5 | -1.5 | -2.5 | A | - | - | - | - | - | Glycopeptide | Antineoplastic | DNA/RNA |
| CID 135500522 | Calcium Levofolinate | 0 | 0 | 2.75 | Y/N | - | - | - | - | - | Tetrahydrofolic acid | Detoxifying agent | Serine hydroxymethyltransferase |
| CID 6915944 | Cefdinir | 0 | 3.5 | 9.5 | Y | 0.5-4 | 0.5-4 | - | - | - | Cephalosporin | Antibiotic | PBP 2-3 |
| CID 44187 | Cefoperazone | 0 | 2.5 | 6.5 | Y | - | - | - | - | - | Cephalosporin | Antibiotic | PBP 1A-B, 2 |
| CID 38103 | Cefradine | 0 | 7.95 | 9.2 | Y | 0.5-4 | 0.5-4 |  |  |  | Cephalosporin | Antibiotic | PBP 1A |
| CID 54682468 | Chlortetracycline HCl | 4.55 | 1.5 | 8.15 | Y | - | - | - | - | - | Aminocyclitol glycoside | Antibiotic | 30S ribosomal |
| CID 71334 | Danofloxacin Mesylate | 7.95 | 0.3 | -0.55 | Y/N | - | - | - | - | - | Fluoroquinolone | Antibiotic | DNA gyrase |
| CID 54680690 | Demeclocycline HCl | 3 | 3.5 | 6.5 | Y | - | - | - | - | - | Tetracycline | Antibiotic | 30S ribosomal |
| CID 21653 | Dihydrostreptomycin sulfate | 6.6 | -6.6 | -0.8 | A | 1 | 0.5-4 |  |  |  | Aminocyclitol glycoside | Antibiotic | 30S ribosomal |
| CID 3229 | Enoxacin | 0 | 5.5 | 5.5 | Y | - | - | - | - | - | Fluoroquinolone | Antibiotic | DNA gyrase |
| CID 71188 | Enrofloxacin | 1.5 | 7.5 | 5 | Y | 0.5-4 | 0.5-4 |  |  |  | Fluoroquinolone | Antibiotic | DNA gyrase |
| CID 5323714 | Furazolidone | 2 | -2 | -2 | A | 1 | 0.75 |  |  |  | Nitrofuran | Antibiotic | DNA |
| CID 5379 | Gatifloxacin | 8.5 | -8.5 | -0.5 | A | - | - | - | - | - | Fluoroquinolone | Antibiotic | DNA gyrase |
| CID 56842139 | Gentamicin Sulfate | 8.75 | 0.4 | 1.65 | Y/N | - | - | - | - | - | Aminoglycoside | Antibiotic | 30S ribosomal |
| CID 23670359 | Ibandronate sodium | 0 | 0 | 3.2 | Y/N | 0.5-4 | 0.5-4 |  |  |  | Bisphosphonate | Osteoporosis | Farnesyl pyrophosphate synthase |
| CID 6321424 | Ivermectin | 2.5 | -2.5 | -2.5 | A | - | - |  |  |  | Milbemycin | Antiparasitic | Glycine receptor |
| CID 149096 | Levofloxacin hydrate | 6.05 | 0.25 | 1 | Y/N | 0.5-4 | 0.5-4 |  |  |  | Fluoroquinolone | Antibiotic | DNA gyrase |
| CID 149096 | Levofloxacin | 6 | -1.5 | -0.5 | A | 0.5-4 | 0.5-4 |  |  |  | Fluoroquinolone | Antibiotic | DNA gyrase |
| CID 68624 | Lomefloxacin HCl | 7.2 | 1.4 | 2.6 | Y | - | - | - | - | - | Fluoroquinolone | Antibiotic | DNA gyrase |
| CID 60651 | Marbofloxacin | 2.5 | 2.5 | 5 | Y | - | - | - | - | - | Fluoroquinolone | Antibiotic | DNA gyrase |
| CID 54685047 | Methacycline HCl | 0 | 3.4 | 8.4 | Y | 0.5-4 | 0.5-4 |  |  |  | Tetracycline | Antibiotic | 30S ribosomal |
| CID 4410 | Nadifloxacin | 0 | 3 | 0 | Y/N | - | - | - | - | - | Fluoroquinolone | Antibiotic | DNA gyrase |
| CID 62115 | Netilmicin Sulfate | 11.45 | -1.775 | 2.45 | Y/N | - | - | - | - | - | Aminoglycoside | Antibiotic | 30S ribosomal |
| CID 4539 | Norfloxacin | 7 | 3 | 1.5 | Y | - | - | - | - | - | Fluoroquinolone | Antibiotic | DNA gyrase |
| CID 4583 | Ofloxacin | 2.5 | 5.5 | 4 | Y | 0.5-4 | 0.5-4 | - | - | - | Fluoroquinolone | Antibiotic | DNA gyrase |
| CID 11478676 | Palbociclib (PD0332991) Isethionate | 0 | 4.85 | 3.45 | Y | - | - | - | - | - | Pyridinylpiperazine | Antineoplastic | CDK4 & CDK6 |
| CID 119525 | Pefloxacin Mesylate | 10.5 | 0.5 | 3.5 | Y | - | - | - | - | - | Fluoroquinolone | Antibiotic | DNA gyrase |
| CID 33630 | Penfluridol | 4 | 0 | 1.5 | Y/N | 0.5-4 | 0.5-4 |  |  |  | Diphenylmethane | Antipsychotic | Dopamine receptors |
| CID 5702105 | Polymyxin B sulphate | 13.55 | -1.1 | 0.3 | Y/N | - | - | - | - | - | [Polymyxin](https://pubchem.ncbi.nlm.nih.gov/compound/polymyxin%20B) | Antibiotic | Lipopolysaccharide layer |
| CID 56207 | Sarafloxacin HCl | 3 | -0.5 | 3 | Y/N | - | - | - | - | - | Fluoroquinolone | Antibiotic | DNA gyrase |
| CID 439243 | Sisomicin sulfate | 10.75 | 0.45 | 3.2 | Y/N | 0.5-4 | 0.5-4 | - | - | - | Aminoglycoside | Antibiotic | 30S ribosomal |
| CID 6918203 | Sitafloxacin Hydrate | 16 | 1 | -2.5 | Y/N | - | - | - | - | - | Fluoroquinolone | Antibiotic | DNA gyrase |
| CID 60464 | Sparfloxacin | 14 | 3 | 4.5 | Y | - | - | - | - | - | Fluoroquinolone | Antibiotic | DNA gyrase |
| CID 36294 | Tobramycin | 4.4 | 1.15 | 1.7 | Y | 0,75 | 0,625 |  |  | * | Aminoglycoside | Antibiotic | 30S ribosomal |
| CID 5564 | Triclosan | 9.5 | 0.15 | -0.3 | Y/N | 0.5-4 | 0.5-4 |  |  |  | Diphenylether | Antiseptic | ENR enzyme |

| ***COLISTIN HITS*** | | **sHTSS** | | | |  |  |  |  |  |  |  |  |
| --- | --- | --- | --- | --- | --- | --- | --- | --- | --- | --- | --- | --- | --- |
|  | |  | **Δ zones of inhibition (mm) ^b^** | |  | **CBA^e^** | | **TKA^f^** | | |  |  |  |
| **PubChem #^a^** | **FDA compound** | **Ø^c^** | **CST 1/256xMIC** | **CST 1/128xMIC** | **Interaction^d^** | **FICI** | **FBCI** | **8h** | **24h** | **48h** | **Chemical classification^g^** | **Therapeutic use** | **Target** |
| CID 2123 | Altretamine | 1.5 | -1.5 | -1.5 | A | - | - | - | - | - | Dialkylarylamine | Antineoplastic | DNA |
| CID 38351 | Amikacin disulfate | 0 | 3.5 | 1.9 | Y | 0.32 | 0.19 |  |  |  | Aminoglycoside | Antibiotic | 30S ribosome |
| CID 16218899 | Amikacin hydrate | 0 | 1.85 | 0 | Y/N | 0.32 | 0.19 |  |  |  | Aminoglycoside | Antibiotic | 30S ribosomal |
| CID 447043 | Azithromycin | 0 | 4.5 | 6.5 | Y | 0.14 | 0.14 | * | * | * | Macrolide | Antibiotic | 50S ribosomal |
| CID 5742832 | Aztreonam | 11 | 2 | 3 | Y | 0.38 | 0.38 |  |  |  | Monobactam | Antibiotic | PBP 3 |
| CID 44599207 | BAF312 (Siponimod) | 0 | 0 | 4.5 | Y/N | - | - | - | - | - | Trifluoromethylbenzene | Multiple sclerosis | S1P receptor |
| CID 72466 | Bleomycin Sulfate | 9.5 | -1 | -6 | A | 0.16 | 0.16 |  | * | * | Glycopeptide | Antineoplastic | DNA/RNA |
| CID 135500522 | Calcium Levofolinate | 0 | 1.5 | 0 | Y/N | - | - | - | - | - | Tetrahydrofolic acid | Detoxifying agent | Serine hydroxymethyltransferase |
| CID 6915944 | Cefdinir | 0 | 5.5 | 0 | Y/N | 1 | 0.63 | - | - | - | Cephalosporin | Antibiotic | PBP 2-3 |
| CID 44187 | Cefoperazone | 0 | 6.5 | 0 | Y/N | - | - | - | - | - | Cephalosporin | Antibiotic | PBP 1A-B, 2 |
| CID 38103 | Cefradine | 0 | 2.85 | 6.4 | Y | 1 | 1 | - | - | - | Cephalosporin | Antibiotic | PBP 1A |
| CID 6536864 | Ceftazidime Pentahydrate | 0 | 3.75 | 0 | Y/N | - | - | - | - | - | Cephalosporin | Antibiotic | PBP 3 |
| CID 137706342 | Ceftriaxone Sodium Trihydrate | 0 | 4.5 | 0 | Y/N | 0.32 | 0.32 |  |  |  | Cephalosporin | Antibiotic | PBP 2B |
| CID 57379345 | Ceritinib (LDK378) | 0 | 0 | 3.5 | Y/N | - | - | - | - | - | Phenylpiperidine | Antineoplastic | Tyrosine kinase receptor |
| CID 71334 | Danofloxacin Mesylate | 7.95 | 2.05 | -7.95 | Y/N | - | - | - | - | - | Fluoroquinolone | Antibiotic | DNA gyrase |
| CID 56205 | Difloxacin HCl | 14 | -2.5 | -1 | A | - | - | - | - | - | Fluoroquinolone | Antibiotic | DNA gyrase |
| CID 21653 | Dihydrostreptomycin sulfate | 6.6 | 1.25 | 0.35 | Y/N | - | - | - | - | - | Aminocyclitol glycoside | Antibiotic | 30S ribosomal |
| CID 71188 | Enrofloxacin | 1.5 | 3.5 | 4 | Y | 0.63 | 0.63 |  |  |  | Fluoroquinolone | Antibiotic | DNA gyrase |
| CID 5323714 | Furazolidone | 5 | -1 | -0.5 | A | - | - | * | * | * | Nitrofuran | Antibiotic | DNA |
| CID 5379 | Gatifloxacin | 8.5 | -7.5 | -8.5 | A | - | - | - | - | - | Fluoroquinolone | Antibiotic | DNA gyrase |
| CID 3476 | Glimepiride | 1.5 | -1.5 | -1.5 | A | - | - | - | - | - | Benzenesulfonamide | Diabetes mellitus | ATP-sensitive potassium channels |
| CID 6321424 | Ivermectin | 2.5 | -2.5 | -2.5 | A | - | - |  |  |  | Milbemycin | Antiparasitic | Glycine receptor |
| CID 149096 | Levofloxacin | 6 | 1 | -2.5 | Y/N | - | - | * | * | * | Fluoroquinolone | Antibiotic | DNA gyrase |
| CID 149096 | Levofloxacin hydrate | 8 | -2.5 | -4.5 | A | - | - | * | * | * | Fluoroquinolone | Antibiotic | DNA gyrase |
| CID 68624 | Lomefloxacin HCl | 7.2 | 0.65 | -7.2 | Y/N | - | - | - | - | - | Fluoroquinolone | Antibiotic | DNA gyrase |
| CID 60651 | Marbofloxacin | 2.5 | 0.5 | -0.5 | Y/N | - | - | - | - | - | Fluoroquinolone | Antibiotic | DNA gyrase |
| CID 65329 | Mefloquine HCl | 0 | 1.5 | 0 | Y/N | - | - | - | - | - | Quinoline | Antimalarial | 80S ribosomal |
| CID 101526 | Moxifloxacin HCl | 3.5 | 2 | -1 | Y/N | - | - | - | - | - | Fluoroquinolone | Antibiotic | DNA gyrase |
| CID 62115 | Netilmicin Sulfate | 11.45 | -0.1 | 0.35 | Y/N | - | - | - | - | - | Aminoglycoside | Antibiotic | 30S ribosome |
| CID 4539 | Norfloxacin | 7 | 0 | 0.5 | Y/N | - | - | - | - | - | Fluoroquinolone | Antibiotic | DNA gyrase |
| CID 4583 | Ofloxacin | 2.5 | 3.5 | 4 | Y | 0.63 | 0.63 | - | - | - | Fluoroquinolone | Antibiotic | DNA gyrase |
| CID 11478676 | Palbociclib (PD0332991) Isethionate | 0 | 4.95 | 6.2 | Y | - | - | - | - | - | Pyridinylpiperazine | Antineoplastic | CDK4 & CDK6 |
| CID 119525 | Pefloxacin Mesylate | 10.5 | -2 | 0.5 | Y/N | - | - | - | - | - | Fluoroquinolone | Antibiotic | DNA gyrase |
| CID 16362 | Pimozide | 0 | 2 | 0 | Y/N | - | - | - | - | - | Diphenylbutylpiperidine | Antipsychotic | Dopamine receptors |
| CID 5702105 | Polymyxin B sulphate | 13.55 | -1.35 | -0.65 | A | - | - | - | - | - | [Polymyxin](https://pubchem.ncbi.nlm.nih.gov/compound/polymyxin%20B) | Antibiotic | Lipopolysaccharide layer |
| CID 56207 | Sarafloxacin HCl | 3 | 5 | 0.5 | Y | - | - | - | - | - | Fluoroquinolone | Antibiotic | DNA gyrase |
| CID 439243 | Sisomicin sulfate | 10.75 | 0.15 | 2.05 | Y/N | 0.5 | 0.38 | - | - | - | Aminoglycoside | Antibiotic | 30S ribosomal |
| CID 6918203 | Sitafloxacin Hydrate | 16 | -2.5 | -0.5 | A | - | - | - | - | - | Fluoroquinolone | Antibiotic | DNA gyrase |
| CID 60464 | Sparfloxacin | 14 | -2 | 4 | Y/N | - | - | - | - | - | Fluoroquinolone | Antibiotic | DNA gyrase |
| CID 36294 | Tobramycin | 4.4 | 1.25 | 1.05 | Y | 0.32 | 0.38 |  |  |  | Aminoglycoside | Antibiotic | 30S ribosome |
| CID 5564 | Triclosan | 9 | 0 | 2 | Y/N | 0.56 | 0.56 |  | * | * | Diphenylether | Antiseptic | ENR enzyme |

| ***FOSFOMYCIN HITS*** | | **sHTSS** | | | |  |  |  |  |  |  |  |  |
| --- | --- | --- | --- | --- | --- | --- | --- | --- | --- | --- | --- | --- | --- |
|  | |  | **Δ zones of inhibition (mm) ^b^** | |  | **CBA^e^** | | **TKA^f^** | | |  |  |  |
| **PubChem #^a^** | **FDA compound** | **Ø^c^** | **FOF 1/8 x MIC** | **FOF ¼ x MIC** | **Interaction^d^** | **FICI** | **FBCI** | **8 h** | **24 h** | **48 h** | **Chemical classification^g^** | **Therapeutic use** | **Target** |
| CID 38351 | Amikacin disulfate | 4 | 0 | 6 | Y/N | 0.5-4 | NA |  |  |  | Aminoglycoside | Antibiotic | 30S ribosomal |
| CID 16218899 | Amikacin hydrate | 2.5 | -1 | -0.5 | A | 0.5-4 | NA |  |  |  | Aminoglycoside | Antibiotic | 30S ribosomal |
| CID 447043 | Azithromycin | 0.5 | 1 | 6.5 | Y | 0.75 | NA |  | * | * | Macrolide | Antibiotic | 50S ribosomal |
| CID 3033819 | Azithromycin Dihydrate | 0 | 2.5 | 2 | Y | 0.75 | NA |  | * | * | Macrolide | Antibiotic | 50S ribosomal |
| CID 5742832 | Aztreonam | 5 | -1.5 | 14.5 | Y/N | - | - | - | - | - | Monobactam | Antibiotic | PBP 3 |
| CID 65958 | Balofloxacin | 0 | 0 | 6 | Y/N | - | - | - | - | - | Fluoroquinolone | Antibiotic | DNA gyrase |
| CID 72466 | Bleomycin Sulfate | 9 | 3 | 15 | Y | 0.56 | 0.56 |  | * | * | Glycopeptide | Antineoplastic | DNA/RNA |
| CID 6915944 | Cefdinir | 7 | 3 | 13 | Y | 0.75 | NA |  |  |  | Cephalosporin | Antibiotic | PBP 2-3 |
| CID 11954009 | Cefmenoxime hydrochloride | 0 | 0 | 9 | Y/N | - | - | - | - | - | Cephalosporin | Antibiotic | PBP 1A |
| CID 44187 | Cefoperazone | 2.5 | 0.5 | 3 | Y | 0.75 | NA | - | - | - | Cephalosporin | Antibiotic | PBP 1A-B, 2 |
| CID 10695961 | Cefotaxime sodium | 0 | 0 | 11 | Y/N | - | - | - | - | - | Cephalosporin | Antibiotic | PBP 1-3 |
| CID 38103 | Cefradine | 3 | -3 | -3 | A | - | - | - | - | - | Cephalosporin | Antibiotic | PBP 1A |
| CID 6536864 | Ceftazidime Pentahydrate | 0 | 0 | 8.5 | Y/N | 0.625 | 0.625 | - | - | - | Cephalosporin | Antibiotic | PBP 3 |
| CID 9937686 | Ceftiofur HCl | 3 | 0 | 15 | Y/N | - | - | - | - | - | Cephalosporin | Antibiotic | PBP |
| CID 137706342 | Ceftriaxone Sodium Trihydrate | 0 | 0 | 4 | Y/N | 1 | NA |  |  | * | Cephalosporin | Antibiotic | PBP 2B |
| CID 1117 | Colistin Sulfate | 11 | -1 | 1 | Y/N | - | - | - | - | - | [Polymyxin](https://pubchem.ncbi.nlm.nih.gov/compound/polymyxin%20B) | Antibiotic | Lipopolysaccharide layer |
| CID 71334 | Danofloxacin Mesylate | 5.5 | 0 | 3.5 | Y/N | - | - | - | - | - | Fluoroquinolone | Antibiotic | DNA gyrase |
| CID 56205 | Difloxacin HCl | 10 | -1 | -3 | A | - | - | - | - | - | Fluoroquinolone | Antibiotic | DNA gyrase |
| CID 21653 | Dihydrostreptomycin sulfate | 5.5 | -1 | -0.5 | A | - | - | - | - | - | Aminocyclitol glycoside | Antibiotic | 30S ribosomal |
| CID 636377 | Doripenem Hydrate | 0 | 0 | 5.5 | Y/N | 1 | NA |  |  | * | Carbapenem | Antibiotic | PBP 1A, 1B, 2-3 |
| CID 54685920 | Doxycycline HCl | 2 | 0 | 3.5 | Y/N | 0.5-4 | NA |  |  |  | Tetracycline | Antibiotic | 30S ribosomal |
| CID 3229 | Enoxacin | 2 | 2 | 1 | Y | - | - | - | - | - | Fluoroquinolone | Antibiotic | DNA gyrase |
| CID 71188 | Enrofloxacin | 8 | 1.5 | 2.5 | Y | 0.5-4 | NA |  |  |  | Fluoroquinolone | Antibiotic | DNA gyrase |
| CID 3374 | Flumequine | 2.5 | 2.5 | 5.5 | Y | 0.5-4 | NA | - | - | - | Fluoroquinolone | Antibiotic | DNA gyrase |
| CID 5323714 | Furazolidone | 0 | 1 | 1.5 | Y | 0.5-4 | NA |  |  |  | Nitrofuran | Antibiotic | DNA |
| CID 56842139 | Gentamicin Sulfate | 2.5 | -0.5 | -2.5 | A | - | - | - | - | - | Aminoglycoside | Antibiotic | 30S ribosomal |
| CID 149096 | Levofloxacin | 5 | 1 | 3.5 | Y | 0.5-4 | NA | - | - | - | Fluoroquinolone | Antibiotic | DNA gyrase |
| CID 149096 | Levofloxacin | 6.5 | -2 | -2.5 | A | 0.5-4 | NA | - | - | - | Fluoroquinolone | Antibiotic | DNA gyrase |
| CID 68624 | Lomefloxacin HCl | 0 | 0 | 3 | Y/N | 0.38 | 0.37 |  |  | * | Fluoroquinolone | Antibiotic | DNA gyrase |
| CID 60651 | Marbofloxacin | 7.5 | -2.5 | -1.5 | A | - | - | - | - | - | Fluoroquinolone | Antibiotic | DNA gyrase |
| CID 54685047 | Methacycline HCl | 4 | -3 | -4 | A | - | - | - | - | - | Tetracycline | Antibiotic | 30S ribosomal |
| CID 54685925 | Minocycline HCl | 3.5 | -3.5 | 2.5 | Y/N | - | - | - | - | - | Tetracycline | Antibiotic | 30S ribosomal |
| CID 441242 | Moxalactam Disodium | 0 | 0 | 7 | Y/N | 1 | NA |  | * | * | Oxacephem | Antibiotic | PBP 1A, 1B, 3 |
| CID 101526 | Moxifloxacin HCl | 5 | 1.5 | 2.5 | Y | - | - | - | - | - | Fluoroquinolone | Antibiotic | DNA gyrase |
| CID 4410 | Nadifloxacin | 6 | -3 | -2.5 | A | - | - | - | - | - | Fluoroquinolone | Antibiotic | DNA gyrase |
| CID 62115 | Netilmicin Sulfate | 9.5 | -1.5 | 1.5 | Y/N | 1 | NA | - | - | - | Aminoglycoside | Antibiotic | 30S ribosomal |
| CID 4583 | Ofloxacin | 0 | 0 | 3 | Y/N | - | - | - | - | - | Fluoroquinolone | Antibiotic | DNA gyrase |
| CID 11478676 | Palbociclib (PD-0332991) HCl | 0 | 2.5 | 0 | Y/N | - | - | - | - | - | Pyridinylpiperazine | Antineoplastic | CDK4 & CDK6 |
| CID 119525 | Pefloxacin Mesylate | 9 | 2 | 2 | Y | 0.75 | 0.625 | - | - | - | Fluoroquinolone | Antibiotic | DNA gyrase |
| CID 33630 | Penfluridol | 5.5 | -0.5 | -0.5 | A | - | - | - | - | - | Diphenylmethane | Antipsychotic | Dopamine receptors |
| CID 5702105 | Polymyxin B sulphate | 8.75 | 0.25 | 0.75 | Y | - | - | - | - | - | [Polymyxin](https://pubchem.ncbi.nlm.nih.gov/compound/polymyxin%20B) | Antibiotic | Lipopolysaccharide layer |
| CID 135445761 | Pralidoxime chloride | 1.5 | 1 | 3 | Y | 0.5-4 | NA |  |  |  | N-methylpyridinium | Organophosphates antidote | Cholinesterase |
| CID 6436173 | Rifaximin | 0 | 0 | 2.5 | Y/N | 1 | NA |  |  |  | Rifaximins | Antibiotic | RNA polymerase |
| CID 56207 | Sarafloxacin HCl | 4 | 1.5 | 5.5 | Y | - | - | - | - | - | Fluoroquinolone | Antibiotic | DNA gyrase |
| CID 439243 | Sisomicin sulfate | 8 | 1 | 2 | Y | 1 | NA | - | - | - | Aminoglycoside | Antibiotic | 30S ribosomal |
| CID 6918203 | Sitafloxacin Hydrate | 18.5 | -1.5 | -3 | A | - | - | - | - | - | Fluoroquinolone | Antibiotic | DNA gyrase |
| CID 60464 | Sparfloxacin | 13 | 1 | 1 | Y | 0.625 | 0.625 | - | - | - | Fluoroquinolone | Antibiotic | DNA gyrase |
| CID 9892071 | Tebipenem Pivoxil | 1 | -1 | 12.5 | Y/N | - | - | - | - | - | Carbapenem | Antibiotic | Unknown |
| CID 441334 | Terbutaline Sulfate | 0 | 2.5 | 0 | Y/N | 0.5-4 | NA |  |  |  | Resorcinol | Antiasthmatic | β-2 adrenergic receptors |
| CID 36294 | Tobramycin | 3 | -2 | -2 | A | - | - | - | - | - | Aminoglycoside | Antibiotic | 30S ribosomal |
| CID 5564 | Triclosan | 2.5 | 0 | 4 | Y/N | 0.5-4 | NA |  |  |  | Diphenylether | Antiseptic | ENR enzyme |
| CID 5578 | Trimethoprim | 0 | 0 | 4.5 | Y/N | 1 | NA |  |  |  | Anisol | Antibiotic | Dihydrofolate reductase |
| CID 35370 | Zidovudine | 4.5 | 0.5 | 9.5 | Y | 0.5-4 | NA |  | * | * | Nucleoside analogue | Antiretroviral | Reverse transcriptase |

^a^PubChem (http://pubchem.ncbi.nlm.nih.gov/) accession number.

^b^Value resulting from the subtraction of the inhibition zone diameter (Ø, mm) of drug-containing agar plates (tigecycline 0.12-0.25 µg/mL, colistin 0.003-0.007 µg/mL and fosfomycin 16-32 µg/mL) from that of drug-free agar plates. Positive values correlate with an increase in the zone of inhibition in the presence of the Primary Compounds (tigecycline, colistin and fosfomycin), indicating possible synergistic interactions. Negative values indicate possible antagonisms.

^c^Ø: Zone of inhibition of the FDA compounds by themselves against K. pneumoniae. Compounds were pin-spotted on agar plates without Primary Compounds.

^d^Interaction classification was based on the increment of the inhibition zones at the two PCs sub-inhibitory concentrations tested compared to those of the no PC plates. Y, synergy (diameter at MICsub1 & diameter at MICsub2 > diameter Ø); Y/N, likely synergy (diameter at MICsub1 OR diameter at MICsub2 > diameter Ø); A, likely antagonism (diameter at MICsub1 & diameter at MICsub2 < diameterØ).

^e^An FICI or FBCI ≤0.5 indicates synergy.

^f^Green indicates synergy. Yellow indicates no interaction. Synergy was defined as a ≥2 log10 reduction in CFU/mL in the combination compared to the most active compound alone. *, cidality under the limit of detection (50 CFU/mL).

^g^From DrugBank (https://go.drugbank.com/)

TGC, tigecycline, CST, colistin, FOF, fosfomycin; FICI, Fractional Inhibitory Concentration Index; FBCI, Fractional Bactericidal Concentration Index; NA, not available.
